# BioContainers Registry: searching for bioinformatics tools, packages and containers

**DOI:** 10.1101/2020.07.21.187609

**Authors:** Jingwen Bai, Chakradhar Bandla, Jiaxin Guo, Roberto Vera Alvarez, Juan Antonio Vizcaíno, Mingze Bai, Pablo Moreno, Björn A. Grüning, Olivier Sallou, Yasset Perez-Riverol

## Abstract

BioContainers is an open-source project that aims to create, store, and distribute bioinformatics software containers and packages. The BioContainers community has developed a set of guidelines to standardize the software containers including the metadata, versions, licenses, and/or software dependencies. BioContainers supports multiple packaging and containers technologies such as Conda, Docker, and Singularity. Here, we introduce the BioContainers Registry and Restful API to make containerized bioinformatics tools more findable, accessible, interoperable, and reusable (FAIR). BioContainers registry provides a fast and convenient way to find and retrieve bioinformatics tools packages and containers. By doing so, it will increase the use of bioinformatics packages and containers while promoting replicability and reproducibility in research.

## 2 Introduction

The BioContainers community has created a complete ecosystem that enables bioinformatics software to be installed and executed in an isolated and controlled environment [2, 8]. Also, it provides infrastructure and basic guidelines to create and distribute bioinformatics containers focusing especially on omics technologies [1]. By August 2020, BioContainers provides over 9,000 tools, 29,000 software versions, and 130,000 packages and containers. It gives access to containers for multiple technologies including Conda [3], Docker, and Singularity [5].

The FAIR Guiding Principles for scientific data management provide recommendations on how to make research data findable, accessible, interoperable, and reusable (FAIR) [9]. In 2017, Jimenez et al. [4] proposed that the FAIR principles for the software involved: i) Findable: the software should be easy to discover by providing software metadata such as title, general description, publication, license, and versions; ii) Accessible: the link to the code or binaries should be available; iii) Interoperable: the metadata should be exchangeable between major software registries; and iv) Reusable: involving the adoption of a license, helping to define to which extent the community can reuse the tool.

Here, we introduce the BioContainers Registry and Restful API (Application Programming Interface) to make bioinformatics tools more findable, accessible, interoperable, and reusable (FAIR). A new registry web interface enables bioinformaticians and developers to easily search for tools using well-defined metadata. Also, we introduce a Restful API implementation of the GA4GH (Global Alliance for Genomics and Health) tool registry specification (TRS) [6]. Finally, a Restful API command line interface was developed allowing containerized based pipelines the utilization of the BioContainers Registry.

## 3 BioContainers

BioContainers infrastructure builds, stores, and releases packages and containers from three different technologies: Conda, Docker, and Singularity. For every bioinformatics software, a Bioconda recipe can be created and the corresponding Conda package built [8]. The new package will be added to the Bioconda channel and the user will be ready to use it in a virtual environment (Figure 1).

**Figure 1.**
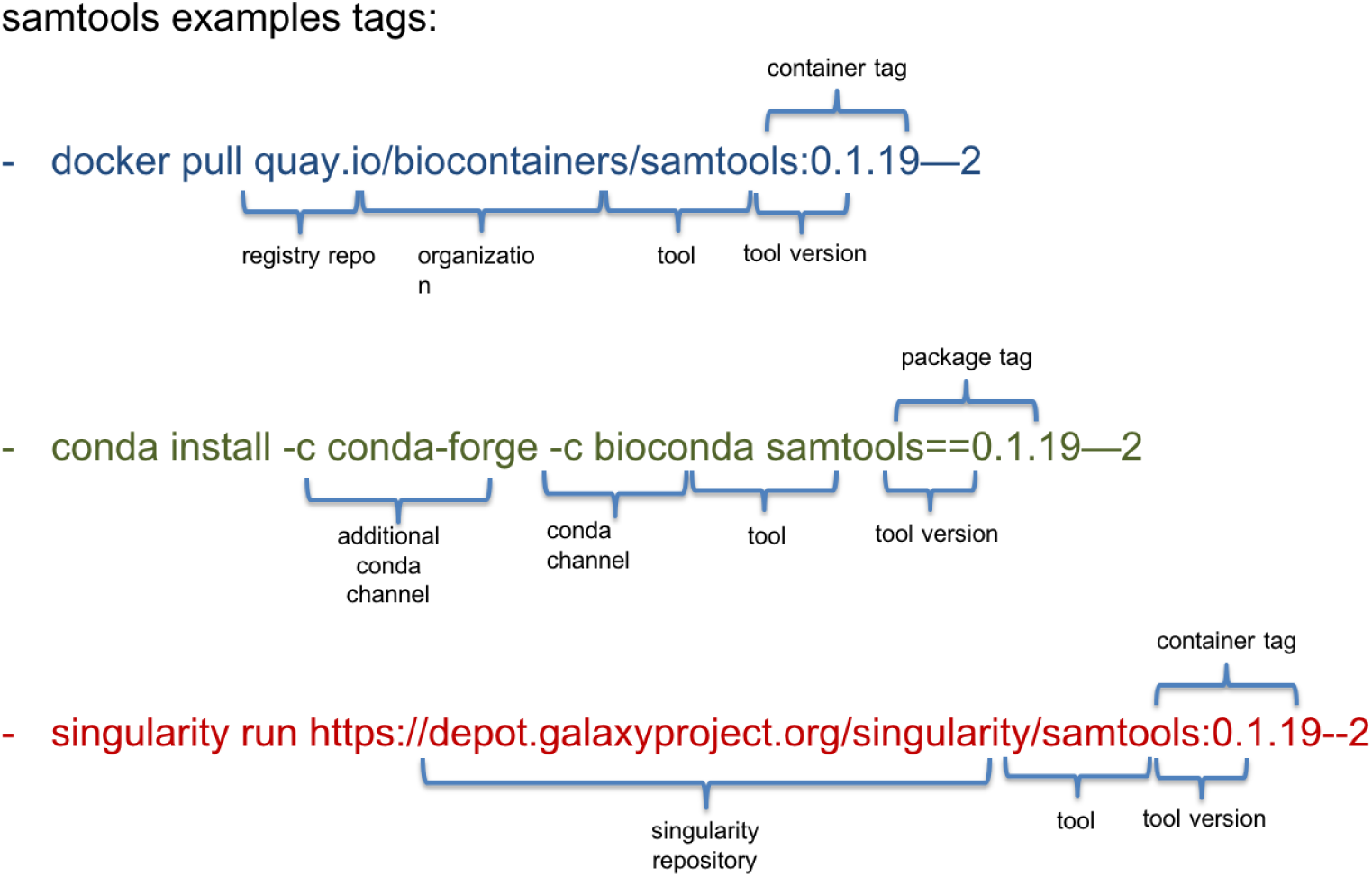
Full tags available for each container technology: Docker, Conda, and Singularity. The full tag is the combination of the repository or registry, the tool name, and the container/package tag.

For example, the Conda package for samtools (Figure 1) can be installed from the following registry (samtools):

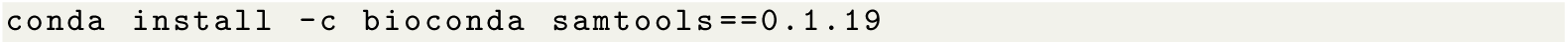

For every Bioconda package, a container is generated automatically and pushed into quay.io. For example, the Docker container for samtools can be retrieved from https://quay.io/repository/biocontainers/samtools (Figure 1). The user can use the samtools containers without needs to install any additional software:

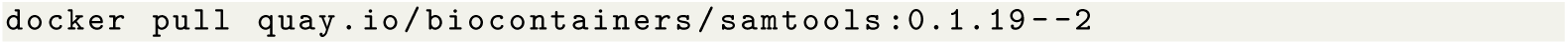

Recently, a third technology, Singularity [5] has emerged to provide more secure containers and High-Performance Computing (HPC) solutions. The BioContainers infrastructure release a singularity image for each docker container in the following repositories: Galaxy depo, Elixir singularity repo. Bioinformaticians and pipelines can use the following command to pull and run Singularity containers:

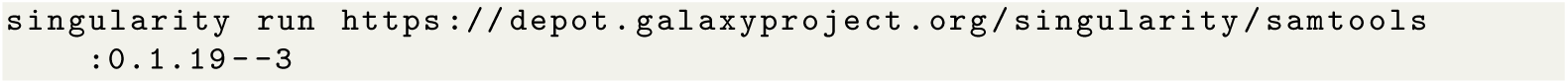

## 4 BioContainers infrastructure and metadata

An important part of the containers and package technologies are the registries and repositories where they are stored. We use multiple registries (Figure 2) for docker (DockerHub, quay.io, Elixir registry), one registry for Conda packages, and two repositories for Singularity images (Galaxy depo, Elixir singularity repo). A continuous integration (CI) system builds the packages in the Bioconda project and pushes the Conda packages to the anaconda registry and to quay.io. For deposition in DockerHub, a Jenkins CI job builds the containers based on the Dockerfile and pushes the images to DockerHub and to the Elixir registry). The use of multiple endpoints for the same technology (e.g. Docker) act as a fail safe, so that when one registry is down, the other registry can serve the containers. This is the case for both Singularity and Docker containers. However, this prompted the need for a centralized metadata registry that enables to search, find and retrieve bioinformatics tools and the corresponding containers.

**Figure 2.**
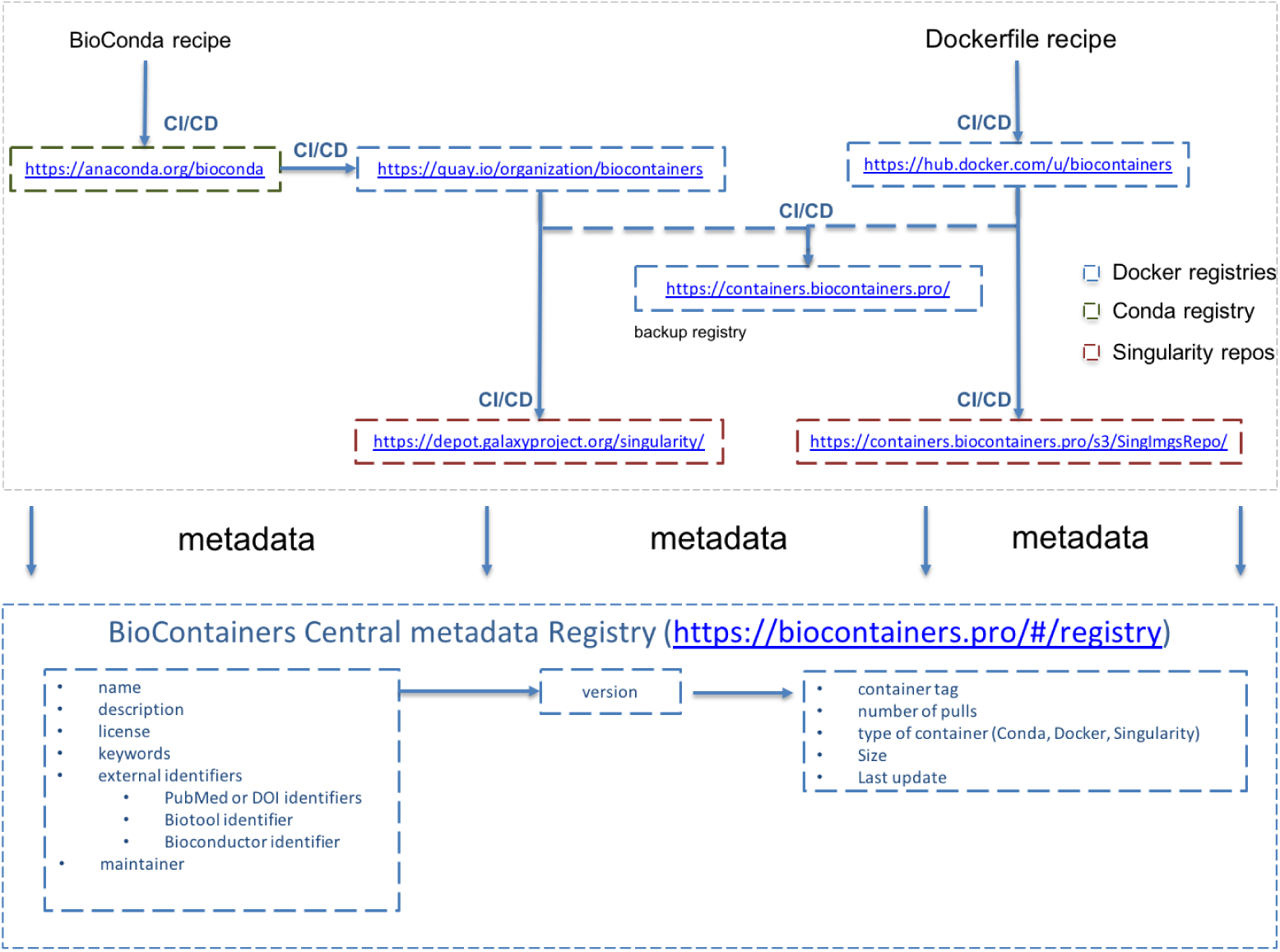
Full URI available for each package technology: Docker, Conda, and Singularity. The full tag is the combination of the repository or registry, the tool name, and the container/package tag.

All the metadata for the tools is stored in a centralized MongoDB database. A set of Python pipelines is used to retrieve the metadata from the Dockerfile and Conda recipes and store them in the database. Also, metadata about the publications and other external identifiers related to the tool are pushed to the database from EuropePMC or biotools. The pipelines, MongoDB database are deployed in a Kubernetes cluster in the ELIXIR compute platform.

### 4.1 Tools and containers metadata

A crucial part of the FAIR principles is the correct annotation of the bioinformatics tools and containers. For each bioinformatics tool, the central registry stores standardized metadata agreed between the Bioconda and BioContainers communities [2]. The ELIXIR tool platform has defined that for every bioinformatics tool the following metadata should be capture (Figure 2) [4]:

The tool metadata includes:

- General metadata: Name, description, license
- Main project URL
- Keywords: transcriptomics, proteomics
- External identifiers such as PubMed accession or DOI (Digital Object Identifier)
- Maintainer
- List of container images

The container metadata includes:

- Container type (Docker, Conda, Singularity)
- Number of pulls (number of times a container has been downloaded/pulled from the original registry)
- Full tag
- Container size
- Date of Last update

The relation between a bioinformatics tool and a container is defined through a set of versions (Figure 2). The database contains two main collections, one for the tool and one for all the versions associated with it. Every version contains a list of containers images with the corresponding metadata.

## 5 BioContainers Registry

The BioContainers Registry provides an easy to use interface for users to search and retrieve the bioinformatics tools and the corresponding containers. The search page contains a search box where users can search using keywords. The search results are then displayed as small boxes, where the name, description, license, and number of downloads are shown. Users can sort using the number of downloads/pulls and filter the results using tool tags and license. Figure 3 shows tool page that describe a selected bioinformatics tool (e.g. samtools). The readme tab (Figure 3) shows the general information about the tool including steps for installation, update, and how to run. In addition, the list of versions, the last update date of the tool, number of downloads, and all the additional identifiers (e.g. PubMed, and bio.tools identifiers) are shown.

**Figure 3.**
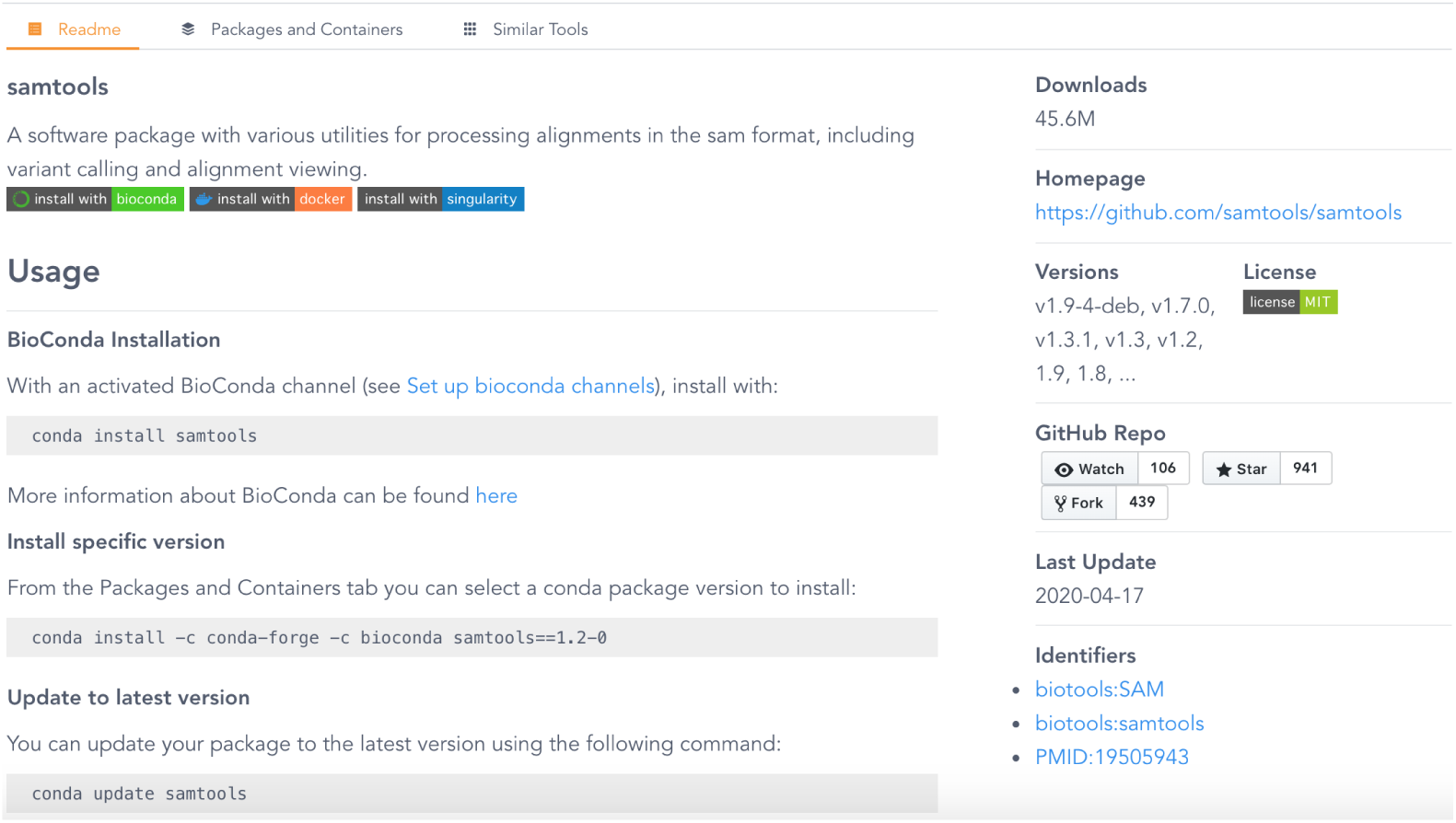
Bioinformatics tool page in BioContainers for samtools

The BioContainers Registry provides for every tool a list of similar tools that can be used to perform the analysis (Figure 4). To enable this, a cosine-similarity algorithm [7] has been implemented using all the metadata available from each tool and container. In the example (Figure 4), a list of tools including sam, perl-bio-samtools, and picard-tools is suggested to the user.

**Figure 4.**
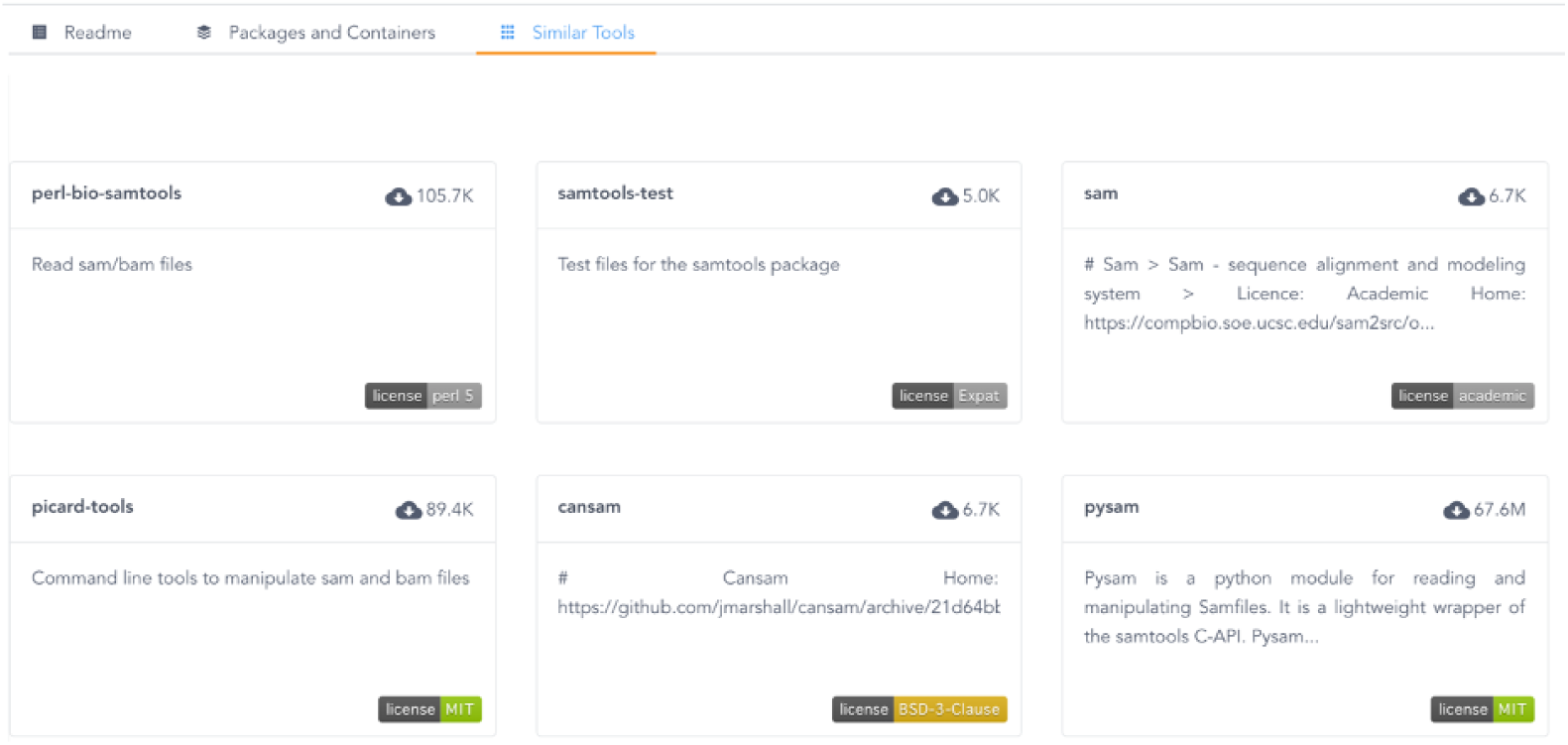
Bioinformatics tool page in BioContainers for samtools

**Figure 5.**
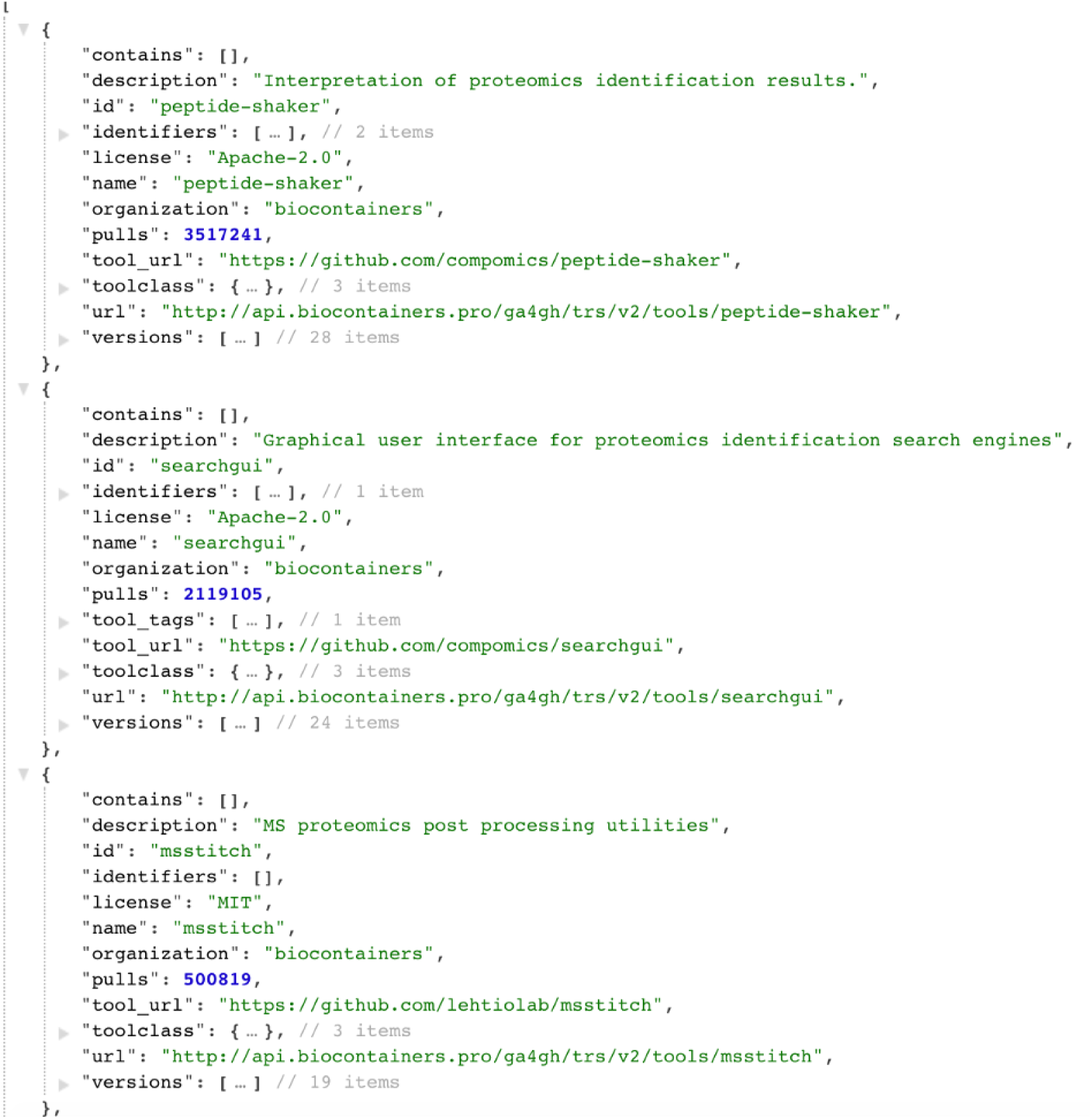
Restful API call for proteomics tools

### 5.1 Restful API

The BioContainers Restful API is an implementation of the TRS standard from the GA4GH consortium [6]. The TRS specification defines an standard API to exchange bioinformatics tools, workflows and containers enabling distribute data processing of omics datasets. The API enables two main functionalities:

- Search for bioinformatics tools and the corresponding containers (/ga4gh/trs/v2/tools): Tools can be searched using the metadata of both the tools and the containers. The API enables to search using specific fields such as id, name, organization, or by using all the metadata fields (e.g. proteomics tools). Also, search results can be refined or filter using different facets (properties) such as license or tool tags (keywords to classify the tools). The following example proteomics and license Apache-2, provides all the tools with proteomics keyword in the metadata and license Apache-2.
- Retrieve the specific tool information and the containers ((/ga4gh/trs/v2/tools/id): All the metadata of the tool, the corresponding versions, and containers are then retrieved (e.g. samtools).

The search results can be sorted by the id, name, organization, description of the tool, the number of downloads, and/or the usage of each tool (Figure 3). The number of downloads and pulls represents the sum of all containers’ downloads and pulls. Sorting the results by the number of downloads enables users to know which tool has been used the most by the community to perform a specific task or which tools are widely used in a particular field (e.g. proteomics, transcriptomics).

## 6 Restful API command line interface

The restful API and web registry interface allows users to easily search and retrieve bioinformatics tools and containers. However, containerized applications are executed in workflows or command-line data analysis pipelines requiring a compatible interface (command line or Python package) to inquire the BioContainers Registry. A python package named bioconda2biocontainer was developed to inquire the registry from the command line interface or through python code as a module. This python package can be used to search Conda packages and versions retrieving the corresponding BioContainers image name. It allows a direct synchronization between Conda environments and containerized analysis pipelines.

The tool can be installed using Conda, Docker or pip:

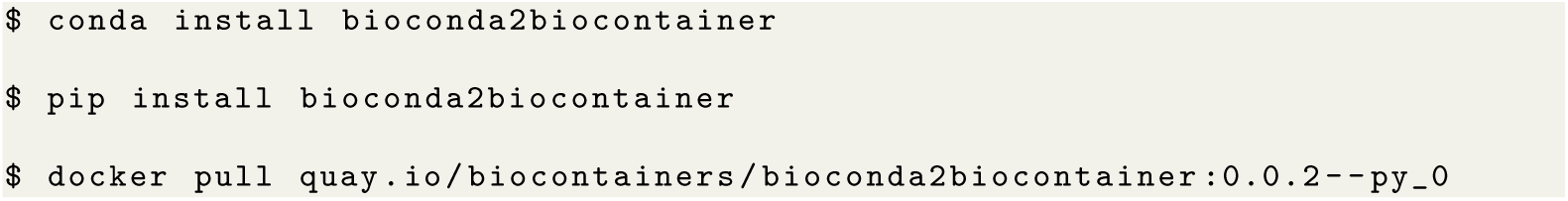

The *biocontainers-search* inquire the BioContainers Registry and return a TAB separated table with Name, Versions (comma separated), Description, License and number of pulls.

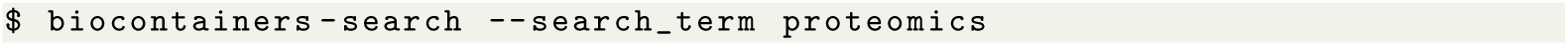

The *–json* parameter can be use output the results in *json* format instead of a tab-delimited output. The name of the tool can be then provided to the *bioconda2biocontainers* tool to find the bioconda, docker or singularity containers available for the tool using the following command:

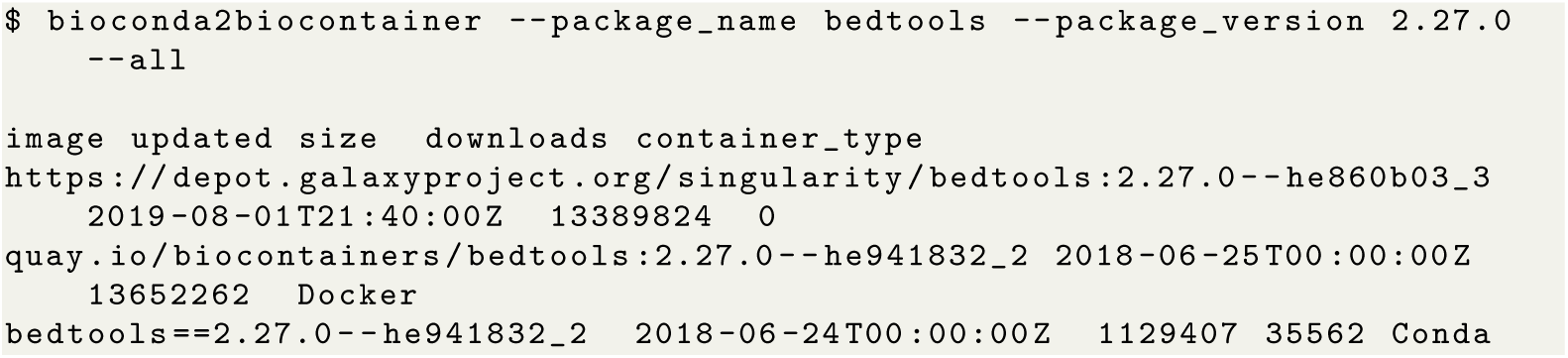

The output of the command is a list of containers for the version of the tool 2.27.0. The images can be sorted by date, size or number of downloads. In addition, the *container type* parameter can be use to retrieve only specific technology containers: Docker, Singularity or Conda.

The *bioconda2biocontainer* package can be used to pull the latest BioContainers Docker image for a Conda package and version as it is shown next:

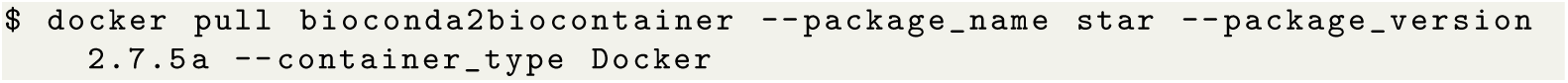

## 7 Conclusion

BioContainers is a growing community with multiple on-going projects, including the maintenance and creation of new software containers for bioinformatics tools, the implementation of improvements in the findability of the tools, and facilitating their re-usability. Recently, a new architecture has been developed to guarantee multiple registries for each container which enables high availability for all the stored BioContainers. Here, we have introduced a new Restful API and web application that can be used to search for bioinformatics tools and its corresponding packages and containers. Researchers can find which are the most downloaded and used tools for a particular task, in addition to having access to all versions of a specific tool. The Restful API is an implementation of the GA4HG standard which enables compatibility with other resources and initiatives like Dockstore.

8

## Abbreviations

API: Application Programming Interface;
FAIR: Findable, Accessible, Interoperable and Reusable;
GA4GH: Global Alliance for Genomics and Health;
HPC: High-Performance Computing;
TRS: Tool Registry Service

## 11 Authors’ contributions

YPR, design the general architecture for the Biocontainer Registry. JB, YPR developed the BioContainer registry Web interface. CB and YPR developed the back-end pipelines and the Restful API. JG and MG contributed with the annotations of the tools. RVA developed the Restful API client. BAG developed the BioConda and Singularity deployment infrastructures and OS and PM implemented the Dockerfile deployment infrastructure. YPR wrote the manuscript with the contributions of PM, JAV and RVA. All authors contribute with the manuscript writing process, read and approved all versions of the manuscript.

## 12 Acknowledgements

All authors would like to acknowledge everyone involved in the Bioconda and BioContainers communities.

